# Dosage responses of aneuploid autosomal chromosomes

**DOI:** 10.1101/2025.09.18.677044

**Authors:** Natali Papanicolaou, Sebastian Wettersten, Guilherme Maia, Antonio Lentini, Björn Reinius

**Author notes:** Co-first author, equal contribution (Natali Papanicolaou, Sebastian Wettersten).

## Abstract

Aneuploidy, resulting from segmental or complete chromosomal losses or gains, is generally detrimental to organismal fitness, but is also associated with tumourigenicity. While sex chromosome aneuploidies are well-tolerated due to dedicated allelic dosage compensation mechanisms, the existence of similar regulatory processes for autosomes remains debated. Here, we investigate transcriptional responses to autosomal aneuploidies and find evidence for global dosage compensation across aneuploid chromosomes. Using high-sensitivity, allele-resolved single-cell RNA-seq on monoclonal cell expansions with varying ploidies and degrees of autosomal aneuploidy, we uncover consistent transcriptional compensation upon chromosome losses via increased burst frequency from the remaining allele. This operates in a region-specific manner across both complete and segmental aneuploidies, revealing a previously underappreciated flexibility of dosage response. Complementary proteomics analyses demonstrate additional dosage buffering at the protein level, resulting in near-stoichiometric rebalancing across autosomal aneuploid chromosomes. Analyses of cancer transcriptomes confirms that compensatory mechanisms are active in primary tumours. Our findings reveal extensive dosage compensation as a genome-wide, dynamic, response to gene product imbalance. This mechanism extends beyond sex chromosomes, supports transcriptional homeostasis, and represents a fundamental, evolutionarily conserved mode of transcriptional regulation active across species and in aneuploid cancer cells.

## Introduction

Chromosome copy number alterations, resulting from chromosomal gains or losses, are known to impair organismal fitness, primarily due to disruptions in protein product stoichiometry^1^. This is reflected in the fact that no autosomal monosomies are viable in humans, and only three autosomal trisomies: trisomies 13, 18 and 21, associated with Patau, Edwards and Down syndromes, respectively. All of these conditions lead to severe phenotypic consequences and, in the case of Patau and Edwards syndromes, early life mortality^2^. Despite these detrimental effects, approximately 90% of solid tumours exhibit aneuploidy, with abnormal chromosome numbers affecting up to 25% of the entire genome^3,4^. Beyond individual chromosome alterations, around 30-60% of all tumours display varying degrees of polyploidy^5–7^, often arising from whole-genome endoduplication and associated with high degrees of aneuploidy and worse overall survival^6,8,9^. While aneuploidy compromises normal fitness, it is also broadly connected to aberrant cellular proliferation and tumourigenic potential^10,11^. However, its precise effects on transcriptional homeostasis and the mechanisms by which it contributes to cancer progression remain poorly understood.

In contrast to the detrimental effects of autosomal aneuploidy, sex chromosomes are functionally aneuploid in the heterogametic sex: male (XY) in mammals and female (ZW) in birds and reptiles. Evolutionary pressures have driven the development of dosage compensation mechanisms to mitigate the harmful effects of sex chromosome monosomies. In therian mammals, this is achieved through inactivation of one X chromosome in female cells, equalising X-linked gene dosage between sexes^12^. As a result, both male and female cells retain a single active X chromosome and are effectively monosomic for the X. To resolve the resulting stoichiometric imbalance between the single X chromosome and paired autosomes, mammals upregulate the active X chromosome to restore dosage balance^13^. This mechanism operates in both sexes, and we have previously shown that its magnitude is elastic, tuning to the degree of X-inactivation in female cells during embryonic development. This tuning is mainly driven by increased transcriptional burst frequency^14,15^. The importance of dosage compensation is further underscored by X-chromosome aneuploidies. Turner syndrome (X0), the only viable monosomy in humans, as well as X-chromosome amplifications such as Klinefelter syndrome (XXY) and trisomy X (XXX), are characterised by relatively mild phenotypic effects. In murine embryonic stem cells with Turner syndrome, dosage imbalances are corrected through X-upregulation^15^. Conversely, in cases of X-chromosome amplification, transcriptional silencing of the additional X copies restores dosage balance^16^. These observations underscore both the necessity of dosage rebalancing and the existence of dosage-sensing mechanisms that guide precise X-inactivation and X-upregulation. Although X-upregulation was long considered mammal-specific, we recently demonstrated that the avian Z chromosome undergoes a similar upregulation through increased burst frequency^17^, suggesting that this mechanism may be more ancient and fundamental than previously believed. Nevertheless, whether such dosage compensation mechanisms are exclusive to sex chromosomes or could extend to autosomal aneuploidy remains a largely open question.

While compensatory mechanisms in response to genetic alterations resulting in mutated mRNAs have been observed^18,19^, these are unlikely to act in numeric copy changes. Although certain oncogenes appear to be dose-regulated through complex regulatory networks^20^, other studies have shown that copy number amplifications of non-oncogenes are often expressed at lower-than-expected levels and can be toxic when overexpressed^21^. These findings suggest that dosage sensing and compensatory mechanisms may extend beyond oncogenes. Several studies in aneuploid yeast strains and human cell lines have reported little to no transcriptional-level dosage compensation for autosomal chromosomes^22–25^. However, more recent work has provided evidence for limited autosomal transcriptional compensation under certain conditions^26,27^. These contrasting findings highlight the complexity of dosage regulation and the need for further investigation into the mechanisms that govern transcriptional responses to aneuploidy. Importantly, previous studies have primarily relied on bulk RNA-sequencing approaches, which are significantly confounded by the presence of mosaic aneuploidies in many tumours, and lack the resolution to distinguish between diploid and tetraploid aneuploidies. Bulk methods average gene expression across heterogeneous cell populations, masking cell-specific chromosomal alterations and underestimating the extent of genomic imbalance. Therefore, disentangling the effect of aneuploidy on individual chromosomes and alleles requires both cellular and allelic resolution, as enabled by single-cell RNA-sequencing (scRNA-seq) approaches^28^. Additionally, allelic information is essential for accurately inferring tetraploidy^5^.

Here, we established clonally expanded aneuploid cells with parental-allele resolution, and using allele-resolved high-sensitivity single-cell RNA sequencing (scRNA-seq) for precise RNA molecule quantification of the dosage response. We demonstrate that autosomal chromosome aneuploidies are transcriptionally compensated at the allele level, achieving partial dosage balance, likely supporting prolonged cellular survival. This compensation is primarily driven by elastic transcriptional regulation via modulation of burst frequency, mirroring mechanisms we previously observed in the mammalian X and avian Z chromosomes, and results in substantial rebalancing at the protein level. In summary, our study provides compelling evidence for both transcriptional and translational dosage compensation of individual alleles in aneuploid cells. It also uncovers, for the first time, the dynamic transcriptional kinetics of aneuploid autosomes. These findings challenge the prevailing notion that such mechanisms are merely functional by-products of sex chromosome evolution and instead indicating more fundamental and conserved molecular dosage compensation processes in the cell.

## Results

### Autosomal chromosomes are dosage compensated in aneuploidy

To enable high-resolution mapping of allelic expression, we derived mouse primary fibroblasts of C57BL/6 × CAST/EiJ F1 hybrid cross. These cells carry a high density of parent-specific single-nucleotide variants (SNVs), allowing allelic resolution in transcriptional readouts. A subset of cells was expanded to allow spontaneous aneuploidy to arise, while another subset was treated with a low dose of Mitomycin C to induce DNA damage, promoting the emergence of cells with varying degrees of aneuploidy and tetraploidy (**Methods**), both spontaneous and induced. Monoclonal expansion was performed from single seeded cells using an automated system in 96-well culturing plates (**Methods**), and 81 clones were selected for screening for allelic imbalances using RNA-seq to identify aneuploid candidates (**Figure 1a** and **Supplementary Fig. 1**). Chromosomal copy number alterations in selected clones were validated by DNA-seq (**Figure 1b**), and ploidy status was confirmed using allelic-specific ratio information (**Methods**), revealing both diploid (2N) and tetraploid (4N) clones. As expected, tetraploid clones exhibited substantially higher aneuploidy scores compared to diploid clones (median 10.5 vs 2.0). This approach enabled the generation of a high-throughput library of clonal aneuploid cell sublines. Importantly, this clonal expansion strategy ensured a high number of genetically near-identical cells, allowing robust inference of individual genetic alterations.

**Figure 1.**
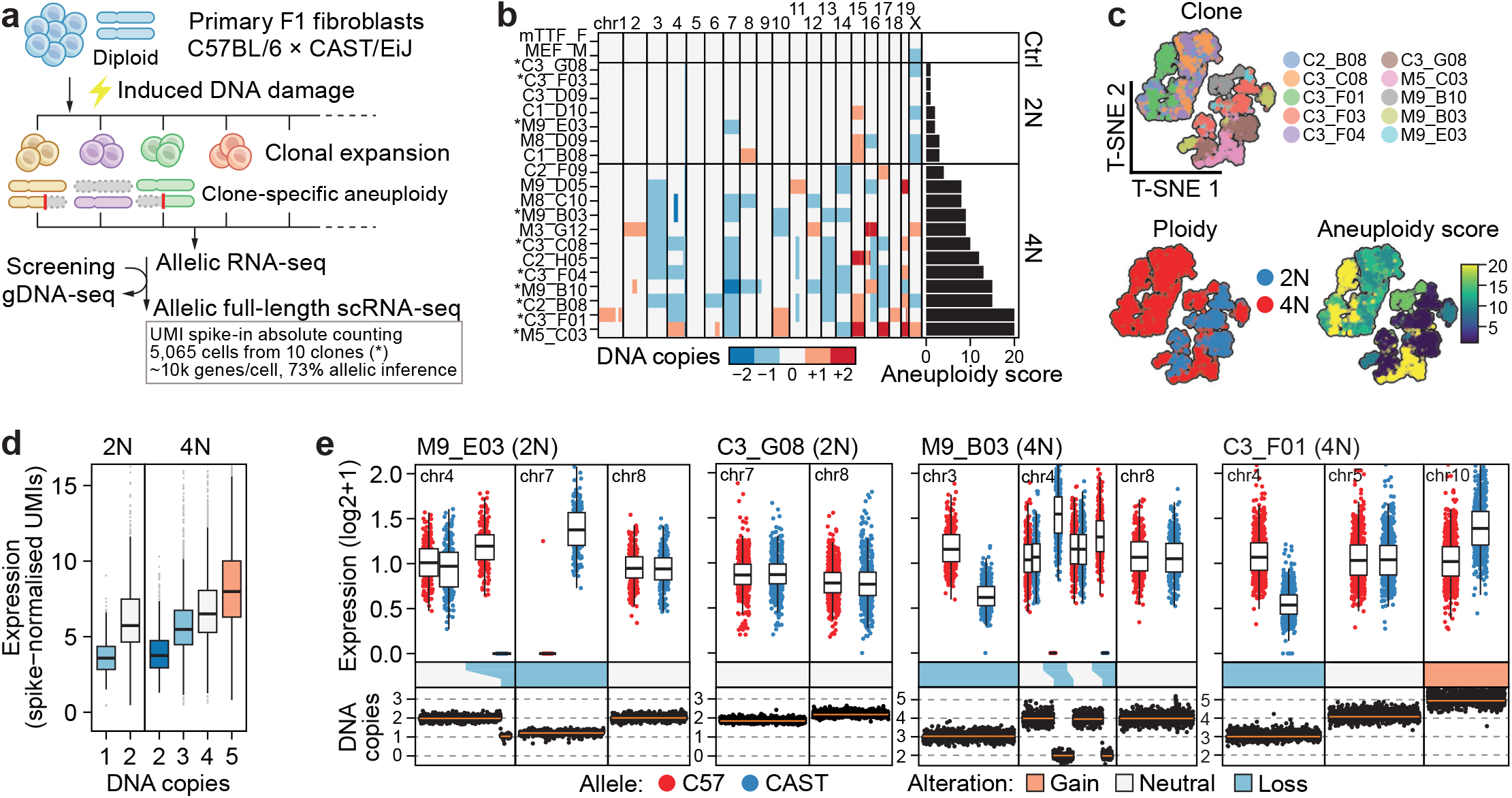
Transcriptional compensation of autosomal aneuploidy. **a**, Experimental design. F1 generation fibroblasts derived from C57BL6/JxCAST/Eij mouse crosseswere treated with Mitomycin C to induce DNA damage. Treated and untreated fibroblasts were then clonally expanded, and colonies were screened for chromosomal abnormalities by allelic bulk RNA-seq. Selected clones were further genotyped using DNA-seq and expanded for allele-resolved full-length single-cell RNA-seq with UMI spike-ins. **b**, Heatmap of aneuploidy score across control, diploid (2N) aneuploid and tetraploid (4N) aneuploid clones based on number of chromosome copies. Untreated clones are labelled “C”, and Mitomycin-C treated clones are labelled “M”. Asterisks (*) indicate clones used for allele-resolved scRNA-seq. Aneuploidy score is defined based on the number of aneuploid chromosomal copies per clone. **c**, Top: t-SNE clustering of aneuploid clones. Bottom left: t-SNE clustering of aneuploid clones based on scRNA-seq coloured by ploidy, with diploid (2N) clones shown in blue and tetraploid clones (4N) shown in red. Bottom right: t-SNE clustering of aneuploid clones based on scRNA-seq gradient coloured by aneuploidy score. **d**, Boxplots of spike-in normalized UMI expression by number of DNA copies for diploid (2N) and tetraploid (4N) aneuploid clones. Data shown as median, first and third quartiles and 1.5x interquartile range. Number of genes per group are indicated. **e**, Boxplots of spike-in normalized allelic expression (log2 +1) for representative diploid (2N; M9_E03, C3_G08) and tetraploid (4N; M9_B03, C3_F01) clones, shown for selected chromosomes. C57BL/6J allele in red; CAST/EiJ allele in blue. At the bottom: matched scatterplots of DNA copies based on gDNA-sequencing data, with copy losses marked in light blue and copy gains marked in orange. Data shown as median, first and third quartiles, and 1.5x interquartile range.

Next, we generated full-transcript-length scRNA-seq libraries across 10 selected clones using a modified version of Smart-seq3xpress^29^ (Xpress-seq), incorporating UMI spike-ins to enable absolute RNA molecule counting, which is critical for accurately capturing transcriptomic responses at count level of original RNA molecules^30^. Crucially, this approach provides superior sensitivity and fidelity at the single-cell level compared to other contemporary scRNA-seq technologies, an essential requirement for the allelic analyses performed. With a total sequencing depth of 7.19 billion reads over 5,065 intact cells (median 224,686 UMIs per cell), we detected a median of 10,889 genes per cell, with over 70% resolvable to individual alleles (**Supplementary Fig. 2a**). Allelic ratios confirmed a high degree of intraclonal similarity and robust clonal identity (**Supplementary Fig. 2b**). Despite distinct genomic alterations, clones exhibited overall transcriptional similarity, segregating primarily by ploidy (**Figure 1c**). Gene set enrichment analysis (GSEA) revealed significant enrichment of GO terms related to cell growth in tetraploid cells, including extracellular matrix organisation, chemotaxis and angiogenesis (FDR=6.44×10^−8^, **Supplementary Fig. 2c**), consistent with the effects of tetraploidisation on cell growth pathways^31^. Globally, gene expression correlated with DNA copy number (Linear Model corrected for cell clone, adjusted R^2^ = 0.34, P < 1×10^−100^), consistent with previous findings^32^. However, transcriptional output showed marked compensation for both chromosomal gains and chromosomal losses using spike-in normalised UMI counts, with an estimated autosomal compensation of approximately 28% (Linear model corrected for cell clone; **Figure 1d** and **Supplementary Fig. 2d**), i.e. partial buffering of dosage effects. Notably, the extent of compensation following the loss of a single chromosome in diploid cells was comparable to the loss of two copies in tetraploid cells, suggesting that dosage compensation is governed by chromosomal stoichiometry rather than absolute chromosome copy numbers. This pattern was consistent across both whole-chromosome and segmental alterations, with transcriptional responses confined to affected regions (**Figure 1e**), supporting a model of elastic dosage regulation that locally mitigates the effects of aneuploidy. To validate these findings, we reanalysed single-cell multi-modal DNA and RNA-seq (DNTR-seq) data from 1,025 human HCT116 cells exposed to etoposide or X-ray irradiation^33^, which induce varying degrees of aneuploidy. As these cells were not clonally expanded, we leveraged the DNA modality to cluster cells based on shared genetic lesions (**Methods**). Consistent with our mouse data, the human HCT116 cells exhibited comparable transcriptome-level dosage compensation for both whole-chromosome and segmental changes **(Supplementary Fig. 3)**. Intriguingly, these autosomal results parallel our previous findings in mouse and chicken sex chromosomes, where gene expression is upregulated in response to genetic monosomy as well as allele-specific removal of transcriptional activity by X-chromosome inactivation^15,17^. Together, these findings suggest the existence of a broader, deeply rooted, and evolutionarily conserved mechanism for dosage sensing and compensation in response to aneuploidy in cells beyond sex chromosomes.

### Increased transcriptional burst frequency buffers autosomal chromosome losses

Our observations raise the question: how might such transcriptional regulation of aneuploidies be achieved? Eukaryotic transcription is inherently stochastic, with gene expression occurring in bursts of activity from each allele^34^. These dynamics can be described by two key parameters: burst frequency (the rate at which transcriptional events occur) and burst size (the average number of RNA molecules produced per transcriptional event). We previously showed that transcriptional upregulation of the mammalian X and avian Z sex chromosomes is driven by increased burst frequency^14,15,17^. To investigate whether similar changes in burst kinetics underlie autosomal dosage responses, we inferred allele- and gene-specific bursting parameters from single-cell Xpress-seq data (**Methods**). Interestingly, transcriptional kinetics in diploid and tetraploid clones occupied a highly similar parameter space for both burst frequency and burst size, suggesting that core principles of burst modulation are maintained despite the presence of additional genome copies (**Supplementary Fig. 4a**). Strikingly, we observed a marked increase in transcriptional burst frequency, without a corresponding change in burst size, on intact alleles following both whole-chromosome and segmental loss of the paired chromosome (**Figure 2a-b** and **Supplementary Fig. 4b**). In contrast, burst kinetics remained largely unchanged in cases of chromosomal gain (**Figure 2a-b** and **Supplementary Fig. 4b**), indicating that the observed dampening of gene expression is not mediated by alterations in transcriptional kinetics. This observation aligns with recent findings suggesting that transcript degradation mechanisms may act on chromosomal amplifications^27^. To validate our findings, we reanalysed an independent allele-resolved scRNA-seq dataset^35^ and identified a clonally expanded primary fibroblast lineage with a subclonal loss of one chromosome 3 allele. This event occurred within approximately seven cell divisions, enabling comparison of two ploidy states separated by only a few divisions (**Supplementary Fig. 5a**). Indeed, consistent with our observations in clonal aneuploid lines, the remaining intact allele exhibited elevated expression and increased burst frequency across the remaining chromosome 3 allele (**Supplementary Fig. 5b-c**).

**Figure 2.**
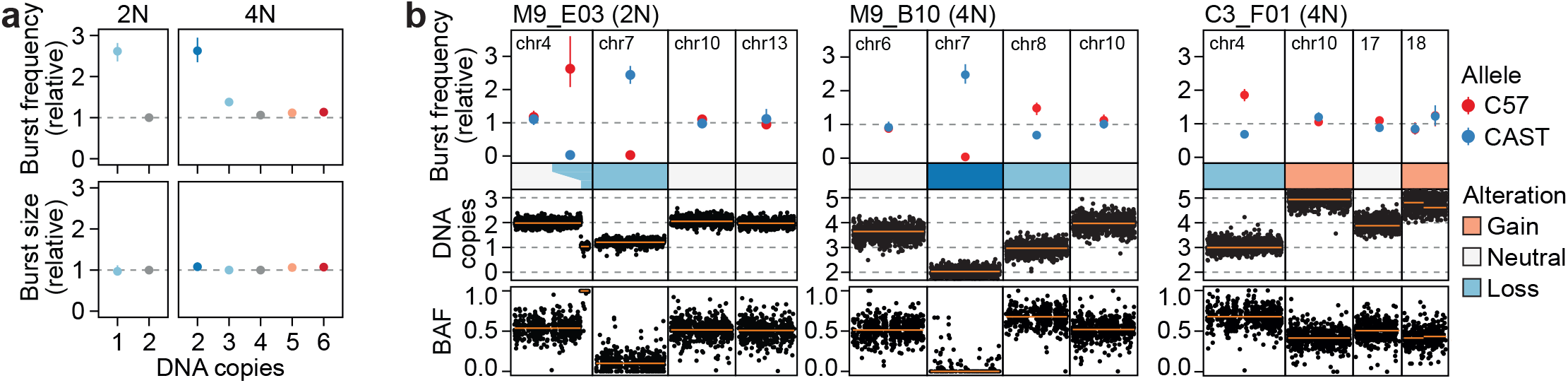
Compensation of autosomal chromosomal losses through increased transcriptional burst frequency. **a**, Top: Median transcription burst frequency across aneuploid chromosomes, relative to median of euploid genes per allele in diploid (2N) and tetraploid (4N) clones. Bottom: Median burst size for aneuploid chromosomes, similarly normalized. Data shown as medians and 95% confidence intervals for the major allele(s) in diploid (1 copy n = 1,192 and 2 copies = 18,164 genes) and tetraploid (2 copies n = 1,121; 3 copies n = 9,335; 4 copies n = 17,511; 5 copies n = 4,977 and 6 copies n = 1,838 genes) clones. **b**, Top: Transcription burst frequency, relative to non-aneuploid chromosomes, shown as median and 95% confidence intervals per allele (C57BL6/J in red, CAST/Eij in blue) for representative diploid (2N) and tetraploid (4N) clones, with selected chromosomes highlighted. Middle: DNA copy numbers per chromosome and clone based on DNA-seq. Bottom: allelic frequency from DNA-seq per chromosome and clone.

The observed increase in transcriptional burst frequency in aneuploid autosomes indicates that this regulatory response is not confined to sex chromosomes. Given the relative tolerance of X-chromosome haploinsufficiency, we sought to directly compare the extent of X-chromosome upregulation with autosomal dosage compensation. Monosomies in diploid cells exhibited transcriptional upregulation ranging from 1.20-to 1.43-fold relative to euploid autosomal alleles. In contrast, X-linked genes were upregulated 1.67-fold **(Supplementary Fig. 6**), consistent with a more efficient dosage compensation on the sex chromosomes. Interestingly, the loss of one X chromosome (X0) in a clone resulted in a similar level of upregulation as observed in XX cells with one active allele due to X-inactivation, further underscoring the elasticity of the upregulation mechanism.

Together, these findings reveal that elastic regulation of transcriptional bursting is a central mechanism driving dosage responses in aneuploidy, including segmental aneuploidies, with a previously unappreciated level of precision. Notably, this response appears to be global, extending dosage compensation beyond sex chromosomes to encompass autosomal regulation.

### Autosomal dosage compensation is further resolved at the protein level and is apparent in cancer

To investigate how transcriptional dosage compensation is reflected at the proteomic level, we performed tandem mass spectrometry on diploid and tetraploid aneuploid clones. Relative protein abundance ratios, normalised against a diploid control, showed a strong positive correlation with chromosomal copy number across both ploidy contexts (adjusted R2 = 0.78, P =6.73×10^−05^; linear model corrected for cell clone). Although RNA and protein expression levels were generally well correlated (**Supplementary Fig. 7a**), protein levels in aneuploid regions were substantially buffered relative to their corresponding RNA levels (**Figure 3a**). Specifically, protein expression was attenuated 12-31% compared to RNA, suggesting the involvement of post-transcriptional regulatory mechanisms in maintaining dosage balance^17–19,27^. Interestingly, chromosomal gains and losses exhibited highly similar linear relationships in their predicted levels of protein-level buffering between diploid and tetraploid cells (**Supplementary Fig. 7b**), indicating that transcriptional and post-transcriptional mechanisms operate in concert to facilitate elastic dosage compensation responses.

**Figure 3.**
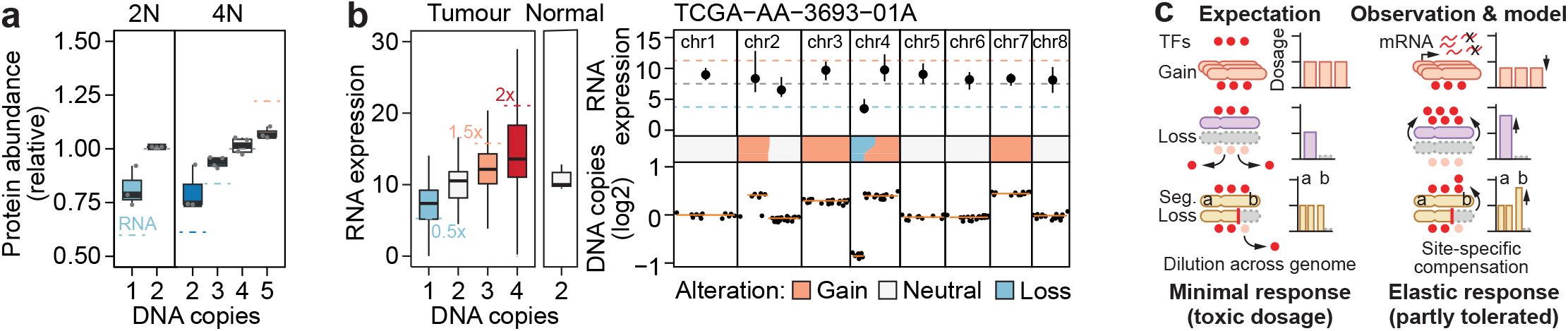
Autosomal dosage compensation extends to the protein level and is conserved in cancer. **a**, Boxplots of protein abundance ratios calculated relative to a diploid control for diploid (2N) and tetraploid (4N) aneuploid clones, based on mass spectrometry. Ratios are grouped by DNA copy number (from DNA-seq). Dashed lines indicate the median RNA expression ratios relative to diploid chromosomal regions. Boxplots are coloured by DNA copy number, and show median, first and third quartiles, and 1.5x interquartile range. **b**, Left: Boxplots of RNA expression levels in tumour (left panel) and normal (right panel) samples versus DNA copy number based on TCGA bulk RNA-seq data. Dashed lines mark expected RNA expression based on ploidy in absence of a dosage compensation response. Right: RNA expression level for chromosomes 1-8 on top, shown as median and 95% confidence intervals. DNA copy numbers for each chromosome shown at the bottom based on matched DNA-seq data for primary tumour sample TCGA-AA-3693-01A. Segmental chromosomal gains highlighted in orange and losses in light blue. **c**, Model of elastic autosomal dosage compensation based on our observations. Left: Aneuploidy caused by chromosomal gains would lead to dilution of the available transcription factors across the additional chromosome copies leading to reduced output from each chromosomal copy. Aneuploidy caused by chromosomal losses would increase the number of transcription factors available, which would be diluted across the genome, leading to minimal dosage compensation from the remaining allele. Right: Based on our observations, chromosomal gains are not compensated at the level of transcription, but at the post-transcriptional and post-translational levels likely through increased degradation of supernumerary transcripts and proteins. However, chromosomal losses are compensated through increased transcriptional burst frequency on the remaining allele, driven partially by an increase in local transcription factor concentrations. This suggests that the additional transcription factors that became available due to chromosomal loss may accumulate at the remaining allele, leading to the elastic dosage compensation we observed.

To assess the relevance of our dosage response findings in the context of cancer aneuploidy, we analysed matched bulk DNA and RNA-seq data from 462 human primary colon adenocarcinoma tumours and 41 normal control samples in The Cancer Genome Atlas (TCGA). We observed a consistent pattern of transcriptional upregulation following chromosomal loss and dampening following chromosomal gain (**Figure 3b**), closely mirroring the trends observed in our clonal aneuploid cell lines and human HCT116 data. For instance, one primary sample (TCGA-AA-3693) exhibited segmental dosage compensation for both chromosomal amplifications and deletions affecting chromosomes 2 and 4 (**Figure 3b**). This observation highlights the elastic nature of the dosage compensation in cancer which appear to act locally on the affected chromosomal segments, similar to recent observations on the X chromosome^36^. Such localised transcriptional regulation may contribute to the cellular tolerance of large-scale aneuploidies in tumours.

## Discussion

In this study, we investigated the existence and mode of dosage compensation in aneuploid autosomal chromosomes using mouse, human, and cancer systems. Leveraging allele-resolved, high-sensitivity scRNA-seq with UMI spike-ins in monoclonally expanded aneuploid cells, we were able to precisely quantify gene expression at allelic resolution across different ploidies. This enabled a detailed characterisation of autosomal gene dosage compensation dynamics, for the first time to our knowledge. Our analyses revealed evidence of widespread elastic transcriptional buffering in response to chromosomal copy number alterations, with compensation proportional to the number of alleles gained or lost. For instance, in diploid cells experiencing complete or partial chromosome loss, the remaining allele exhibited increased transcriptional output of around 20-40%, resulting in substantial dosage rebalancing on a gene-by-gene basis. In contrast, tetraploid cells with a single-copy loss showed a more modest upregulation, underscoring the plasticity of the dosage response and suggesting that compensation is governed by the stoichiometry of gene products rather than absolute copy number changes. These findings were further corroborated in human colorectal adenocarcinoma cells (HCT116) and human primary colon adenocarcinoma tumours, providing compelling evidence for the existence of fundamental dose-compensatory circuits acting on autosomal chromosomes across mammalian species and in cancer.

Using allele-resolved scRNA-seq, we dissected the transcriptional burst kinetics in clonal lineages and found that the transcriptional upregulation following chromosomal loss is primarily driven by increased burst frequency. Mechanistically, this response mirrors known dosage compensation strategies observed in mammalian X-chromosome upregulation and avian Z-chromosome upregulation^14,15,17^, suggesting an ancient and evolutionarily conserved regulatory mechanism for responding to chromosomal loss. Importantly, while chromosome gains were associated with buffered expression levels, the underlying burst kinetics remained largely unchanged. This suggests that, in contrast to chromosomal losses where compensation occurs via increased transcriptional activity, gains are buffered through post-transcriptional mechanisms that modulate mRNA abundance. This finding is consistent with previous findings indicating enhanced degradation of gene products in amplified genomic regions^27,37,38^, highlighting the importance of multiple, complementary layers of dosage regulation in preserving transcriptional homeostasis. Indeed, using high-sensitivity proteomic measurements, we demonstrate that the extensive, yet incomplete, autosomal dosage compensation observed at the transcriptomic level is further consolidated at the proteomic level, resulting in substantial stoichiometric rebalancing. This is consistent with previous proteomic studies finding deviations in protein levels relative to expectations based on DNA copies^27,37,38^.

While dosage-sensitive genes such as MYC have been proposed to rely on complex compensatory mechanisms involving microRNA-transcription factor feedback loops^20^, the majority of genes do not exhibit such sensitivity. This suggests the existence of a more generalised dosage-sensing mechanism. Although its precise nature remains unclear, insights from sex chromosome regulation offer valuable clues. In diploid cells, only one X chromosome remains transcriptionally active, whereas in triploid (XXY) and tetraploid (XXXX, XXXY, XXYY) cells, two X chromosomes are generally active^16,39^. This pattern points to an intrinsic mechanism that maintains stoichiometric balance in relation to cellular ploidy. Under normal expectations, the loss of a chromosomal copy would lead to dispersion of available transcription factors across the genome, resulting in limited compensation for the missing chromosome (**Figure 3c**). Contrary to this expectation, our findings demonstrate a precise, site-specific transcriptional compensation mechanism, driven by increased burst frequency, observed in both diploid and tetraploid cells. This suggests a local enrichment of transcription-driving factor concentration at the remaining allele^40^, potentially facilitated by physical compartmentalisation of transcription and its drivers^41^. Such compartmentalisation may enhance transcription factor retention site-specifically^42^, enabling non-linear transcriptional responses. Similar mechanisms have been described for the MSL2 complex in *Drosophila*, where compartmentalisation and expression upregulation is achieved specifically for the X chromosome despite sharing binding motifs with autosomes^43^. Interestingly, local RNA concentrations have been shown to regulate the formation of condensates and transcriptional bursts^44^, which may explain why burst frequency rather than burst size is modulated to achieve upregulation. Intriguingly, recent data further support the elastic nature of these dosage compensation mechanisms, revealing that the X chromosome can undergo localised transcriptional upregulation in response to CRISPR-induced segmental deletions^36^. While incomplete dosage compensation would typically reduce cellular fitness in normal cells, it may confer a selective advantage in transformed cells that lack replication checkpoints and exhibit replicative immortality. Moreover, because stoichiometric imbalances caused by aneuploidy are less pronounced in tetraploid cells for equivalent chromosomal changes, whole-genome duplications may be favoured as a strategy to mitigate fitness disadvantages.

Together, our data show that cells employ local regulatory strategies that can act with precision at the affected chromosomal regions to preserve transcriptional balance, thereby contributing to cellular resilience when faced with aneuploidy.

## Methods

### Animal housing and ethics statement

Mice were housed in specific pathogen-free at Comparative Medicine Biomedicum (KM-B), Karolinska Institutet, according to Swedish national regulations for laboratory animal work food and water ad libitum, cage enrichment, and 12 h light and dark cycles. All animal experimental procedures were performed in accordance with Karolinska Institutet’s guidelines and approved by the Swedish Board of Agriculture (**permits 17956-18 and 18729-2019** Jordbruksverket).

### Primary mouse fibroblast derivation and culturing

Primary mouse tail tip fibroblasts were isolated from tail explants of 8 week old hybrid CAST/Eij x C57BL6/J mice. Briefly, mice were sacrificed by cervical dislocation and the skin and tail were sterilised with 70% ethanol. The tail was cut, skinned and washed in ice-cold PBS. The tissue was further minced into 5 mm-long pieces and placed in 0.s1% gelatin-coated 10 cm culture dishes with 6 ml of complete media (DMEM – Glutamax [Gibco], 10% heat-inactivated fetal bovine serum (FBS) [Gibco], 100U/ml Penicillin-100μg/ml Streptomycin [Gibco], 1mM sodium pyruvate [Gibco], 1mM non-essential amino acids (NEAA) [Gibco]) and placed at 37°C for 5 days to allow for fibroblasts to migrate out of the explants. After 5 days, the explants were removed and the media was replaced. The following day, the cells were passaged for the first time to allow for even monolayer formation. Briefly, the media was removed, the cells were washed once with PBS and detached using 2 ml TryPLE. Following centrifugation at 300xg for 5 min, the supernatant was removed, the pellet was washed in 2ml PBS and spun down. The final pellet was split 1:2 and plated onto 0.1% gelatin-coated 10 cm dishes containing 8ml of complete mTTF media, and placed at 37°C, 5% CO_2_.

### Treatment with Mitomycin C

Mouse tail tip fibroblasts (mTTFs) were treated with complete media supplemented with 10μM Mitomycin C in PBS for 1h before replacing the treatment with pre-warmed complete media. The cells were incubated at 37°C and 5% CO_2_ for 3 days to recover after mitomycin-C treatment.

### Clonal expansion of fibroblast cells

Single cells from Mitomycin-C-treated and control mTTF cultures were dispensed in 0.1% gelatin-coated 96-well plates containing 100 μl pre-warmed and pre-equilibrated complete media using an UP.SIGHT Single-Cell Dispenser instrument [Cytena] to ensure monoclonal formation of all colonies. Sorted plates were incubated at 37°C, 5% CO_2_ and colony formation was closely monitored in every well of 96-well plates over the next 3 weeks. Every third day, 50% of the media was replaced with fresh complete media until colony size reached an approximate content of about 100 cells, after which media volume was increased to 200 μl. In total, 1,152 individual cells were seeded and 8% of them led to expansions with sufficient proliferation for further use. Once colony-containing wells became confluent, clonal cells were collected from all surviving colonies for cell freezing and mini-bulk Smart-seq2 for colony screening (see section “***Mini-bulk Smart-seq2 library preparation for colony screening”***).

### Mini-bulk Smart-seq2 library preparation for colony screening

#### Clone collection

Clones were grown to confluency in 0.1% gelatin-coated 96-well plates in 200 μl of complete media (DMEM - Glutamax, 10% heat-inactivated fetal bovine serum (FBS) [Gibco], 100U/ml Penicillin-100μg/ml Streptomycin [Gibco], 1mM sodium pyruvate [Gibco], 1mM non-essential amino acids (NEAA) [Gibco]). When confluent (each well containing 20000-40000 cells), the media was removed and the cells thoroughly washed with PBS, before detaching using 100 μl TryPLE and centrifugation at 300 x g for 5 min. Each cell pellet was then resuspended in 1 ml ice-cold PBS, and 10 μl of cell suspension was transferred into 10 μl of ice-cold PBS in a 96-well PCR plate [Armadillo, Thermofisher]. After samples were collected from all clones, the PCR plate was centrifuged for 2 min at 300 x g to pellet the cells and 19 μl of supernatant was carefully removed (leaving 1 μl of supernatant) using a Biomek NXp liquid handling robot [Beckman Coulter].

#### Library preparation

Mini-bulk Smart-seq2 libraries were then prepared as previously described with slight modifications^45^.

#### Cell lysis and reverse transcription

Briefly, cells were lysed by adding 3.5 μl of lysis buffer (containing: 0.1% Triton-X-100 [Sigma Aldrich], 2 mass Units/μl SEQURNA RNase inhibitor^46^ [SEQURNA], 2 mM (each) dNTP mix [Thermofisher], 2 μM SS2 oligo-dT_30_VN [5′-AAGCAGTGGTATCAACGCAGAGTACT30VN-3′; IDT], followed by incubation at 72°C for 3 min and reverse transcription by adding 5.5 μl of reverse transcription mastermix (1x Superscript II first-strand buffer, 5mM betaine [Sigma], 6mM MgCl_2_ [Ambion], 1μM TSO [5′-AAGCAGTGGTATCAACGCAGAGTACATrGrG +G-3′; IDT], 17U/μl Superscript II reverse transcriptase [Thermofisher] to each sample and incubation using the following program: 42°C for 90 min, 10 cycles of [50°C for 2 min and 42°C for 2 min], followed by 70°C for 15 min and 4°C on hold. *PCR pre-amplification*. PCR pre-amplification was performed by adding 15 μl of pre-amplification PCR mastermix (1x KAPA HiFi HotStart Readymix [Roche], 1μM ISPCR oligo [5′-AAGCAGTGGTATCAACGCAGAGT-3’; IDT]) to each sample and incubating using the following thermocycler program: 98°C for 3 min, 16 cycles of [98°C for 20 sec, 67°C for 15 sec, 72°C for 6 min], 72°C for 5 min and 4°C on hold *Bead purification*. The PCR products were purified by adding AMPure XP beads to each sample with a bead:sample ratio of 0.8:1 (20 μl of AMPure XP beads to 25 μl of sample). The samples were incubated at room temperature for 8 min and then placed on a magnetic rack for 5 min. The supernatant was removed, the beads were washed twice with 200 μl of freshly-prepared 80% ethanol, and left to air-dry for 3 min. The cDNA was eluted in 17 μl of nuclease-free water [Ambion]. Library quality was assessed using a Bioanalyzer high sensitivity dsDNA chip and all samples were quantified using the Quantifluor dsDNA system [Promega] according to the manufacturer’s instructions to obtain accurate cDNA concentrations.

#### Library tagmentation, indexing and library amplification

The libraries were normalised to 1 ng/μl and tagmentation was performed by combining 2 ng of cDNA with 18 μl of tagmentation mix containing 10mM TAPS-NaOH [Sigma], 5mM MgCl_2_ [Thermofisher], 8% PEG-8000 and 0.5 μl of in-house produced Tn5 at a concentration of 44.5μM. The samples were incubated at 55°C for 8 min. To strip the Tn5 off the tagmented DNA, 3.5 μl of 0.2% SDS was added to each sample and the mixture was incubated at room temperature for 5 min. The samples were indexed using 1.5 μl of combined, custom-made Nextera i7 and i5 indexes [IDT] and PCR-amplified using 25 μl of PCR mastermix per sample (1x KAPA HiFi PCR buffer, 0.06 mM (each) dNTPs, 1U KAPA HiFi polymerase). The reaction took place as follows: 72°C for 3 min, 95°C for 30 sec, 10 cycles of [95°C for 10 sec, 55°C for 30 sec, 72°C for 30 sec], 72°C for 5 min and 4°C on hold. The libraries were pooled and purified as described above (see section “*Bead purification*”). *Library quality control and sequencing*. The final library pool fragment size was inspected on an Agilent 2100 Bioanalyzer instrument using a High-Sensitivity dsDNA chip [Agilent] and library concentration measured in a Qubit 4.0 fluorometer using the Qubit dsDNA High Sensitivity Assay Kit [Thermofisher]. Sequencing was performed on an Illumina Nextseq 550 instrument using a Nextseq 500/550 High-Output 75 cycle sequencing kit v2.5 [Illumina] using the following settings: Read 1 = 72 cycles, Index 1 = 10 cycles, Index 2 = 10 cycles.

#### Data analysis

Raw Mini-bulk Smart-seq2 data was processed using the zUMIs pipeline^47^ (v2.9.7c). In short, this pipeline filters sample barcodes, the data is aligned with STAR^48^ (v.2.7.2a, options -- limitSjdbInsertNsj 2000000 --clip3pAdapterSeq CTGTCTCTTATACACATCT), reads are assigned to both intron and exon features using featurecounts^49^, barcodes were collapsed by 1 hamming distance, and gene expression was calculated for reads. Allele-level expression was calculated from the reads output by zUMIs using previously described^50^ (github.com/sandberg-lab/Smart-seq3/tree/master/allele_level_expression).

### DNA isolation and preparation of DNA-seq libraries

#### DNA isolation

DNA was extracted from ~ 100000 cell pellets from each clone using Monarch’s genomic DNA purification kit according to the manufacturer’s instructions. Briefly, frozen cell pellets were slowly thawed on ice and resuspended in 100 μl ice-cold PBS. The samples were then treated with 1 μl Proteinase K and 3 μl RNase A and cells were lysed using 100 μl cell lysis buffer, followed by incubation at 56°C for 5 min with agitation at 1400 rpm. Next, 400 μl gDNA binding buffer was added and each sample was thoroughly mixed by pulse vortexing. The lysates were transferred to a gDNA purification column placed in a collection tube and centrifuged for 3 min at 1000 x g and 1 min at maximum speed (12000 x g). The column was transferred to a new collection tube and the column was washed with 500 μl gDNA wash buffer and centrifuged at maximum speed (12000 x g) for 1 min twice. Finally, gDNA was eluted from the column by adding 40 μl pre-warmed gDNA Elution buffer and centrifuging at maximum speed (12000 x g) for 1 min. DNA concentration was measured using a Nanodrop 2000 instrument. To prepare for DNA-seq library preparation, the isolated gDNA was diluted to 1ng/μl.

#### DNA-seq library preparation

For DNA-seq library preparation, 5 ng of gDNA (1ng/μl) were incubated with 15 μl of tagmentation mastermix (10mM TAPS [Sigma], 5mM MgCl_2_ [Thermofisher], 10% dimethylformamide [Sigma], 2.25μM Tn5 [in-house, produced as previously described^51^] at 55°C for 8 minutes. Tn5 was stripped from the tagmented DNA by adding 3.5 μl 0.2% SDS to each sample. The samples were briefly centrifuged and incubated at room temperature for 5 min. Indexing was performed using 2.5 μl of 1 μM pre-mixed Nextera index primers [IDT]. Post-tagmentation PCR was performed by adding 16.5 μl of PCR master mix (1x KAPA HiFi PCR buffer [Roche], 0.6mM (each) dNTPs [Thermofisher], 1U/μl KAPA HiFi polymerase [Roche]) to each sample and incubating using the following thermocycler program: 72°C for 3 min, 95°C for 30 sec, 6 cycles of 95°C for 10 sec, 55°C for 30 sec, 72°C for 30 sec, followed by 72°C for 5 min and 4°C on hold. Double purification of the indexed libraries was performed using 22% PEG magnetic beads prepared as previously described^50^. Briefly, the indexed libraries were combined with 22% PEG beads at a bead:sample ratio of 0.9:1 and the mixture was incubated at room temperature for 8 min. The samples were placed on a magnetic rack for 5 min. Once clear, the supernatant was removed and the beads washed twice with freshly-prepared 80% ethanol. The beads were then left to air-dry for 3 min while placed on the magnetic rack. The samples were then eluted in 30 μl nuclease-free water. The second purification was performed as described above and the purified libraries were eluted in 18 μl of nuclease-free water [Ambion]. Library fragment size was assessed on an Agilent 2100 Bioanalyzer instrument using a Bioanalyzer High Sensitivity dsDNA chip and library concentrations were quantified using Qubit’s high-sensitivity dsDNA quantification kit on a Qubit 4.0 Fluorometer. Libraries were pooled in equimolar amounts and sequenced on a Nextseq 550 instrument using a Nextseq 500/550 High-Output 150 cycle sequencing kit using the following settings: Paired-end, Read 1 = 74, Index 1 = 10, Index 2 = 10, Read 2 = 74).

#### Data analysis

Data was quality and adapter trimmed using fastp^52^ (v0.20.0) and aligned to an N-masked genome using minimap2^53^ (v2.24-r1122, - ax sr). Binned read counts were calculated for 100kb bins using bedtools^54^ (v2.30.0, makewindows -w 100000, multicov -p -q 13) and allele frequencies were calculated for known phased genetic positions using bcftools^55^ (v1.10.2, mpileup -a AD,DP --max-depth 8000, call -mv) and furthered filtered for only variants matching phased allele bases (bcftools isec -n =2 -w 1). Allelic count tables were generated using bcftools (query -f ‘%CHROM %POS %REF %ALT [%AD]\n’). To generate bin-level annotations, mm10 mappability tracks were obtained for k50 multi-read mappability^56^ and averaged per 100kb bin using bedtools (map -c 4 -o mean). Nucleotide frequencies per bin were calculated for the N-masked genome using bedtools (nuc) and bins located within 500kb of large assembly gaps (e.g. centromeric regions, obtained from UCSC mm10 gap table) were identified using bedtools (window -w 500000 -c). Fraction of bins being repetitive was calculated from the UCSC mm10 RepeatMasker table using bedtools (intersect -wao -a bins -b rmsk, map) and awk by overlapping regions and calculating fraction of overlap. Genome bins with >2.5% N bases or average mappability <55% or within 500kb of gaps or being >70% repetitive were excluded and a 5 bin rolling median was applied to smoothen and exclude bins in large excluded regions. Next, read counts were corrected for GC-content and mappability using HMMcopy (correctReadcount, mappability = 0.55, samplesize = 5e3) and normalised per bin against a normal mTTF sample. Genomic copy number segments were identified using a hidden markov model in HMMcopy (setting parameters e=0.9999999999999999 and strength=1e30). Ploidy and cellularity was estimated using ACE^57^ (squaremodel, penalty=1, penploidy=0.5, method=“MAE”) and corrected copy numbers were obtained. Allelic ratios were summarised per 100kb bin using only position with at least 3 reads. Ploidies were further refined and confirmed manually using allele frequencies, where ploidies had to be compatible with median allele frequencies per segment for the majority of chromosomes in each sample. Aneuploidy scores were calculated for autosomes as the sum absolute integer copy number difference between segments against ploidy per sample.

### Full-transcript-length single-cell RNA-seq with UMI spike-ins

#### Library preparation

Full-length single-cell RNA-seq library preparation using the Xpress-seq (v1) method was performed at Xpress Genomics (Stockholm, Sweden). In brief, single cells were sorted using a Sony SH800S instrument into 384-well plates containing lysis buffer, spun down and stored at −80 °C. Upon submitting plates to Xpress Genomics, robotic automated library preparation was performed. Sequencing was performed on the DNBSEQ G400RS platform (MGI Tech) using App-C Sequencing primers.

#### Data analysis

Raw Xpress-seq data was processed using the zUMIs pipeline (v.2.9.7c) as described above in the Mini-bulk Smart-seq2 section with the notable exception that gene expression and allele-level expression were calculated for both reads and UMIs. Spike-in UMI information was extracted for complex molecular spikes from aligned bam files using UMIcountR^58^ using a maximum pattern distance of 2 and corrected using the “adjacency directional” method with a hamming distance of 1 per sample. Spike-ins were filtered for sequences captured in more than 5 barcodes or with more than 100 reads. Cells containing 10% or more spike-in UMI reads were excluded and then filtered for cells with low gene detection (3 MADs lower on log-scale). Spike-in UMI scaling factors were calculated and used for normalisation using scater/scran^59^. Average expression per cell, allele and segment was calculated as 20% trimmed means for genes with a spike-in normalised UMI count > 0. Only segments containing more than 100 detected genes (spike-in normalised UMIs > 0) per clone were used for global analyses. For dimensionality reduction and batch integration, top 500 highly-variable genes (FDR<0.05) were selected in a batch-aware manner and integrated using scVI^60^ (gene_likelihood = ‘nb’) using raw UMI counts to avoid biases related to inherent differences in absolute mRNA counts between ploidies. T-SNE was then calculated on the scVI latent dimensions. GSEA analysis was performed on average gene-wise log2 fold changes for GO BP terms using clusterProfiler and terms with an FDR<1×10^−4^ were kept. Transcriptional burst kinetic inference and analysis was performed using txburst for allelic spike-in normalised UMIs and TPMs, as previously described^15,34^. Genes showing poor inference quality were excluded and data was normalised relative to median per clone. For burst frequency, spike-in normalised TPM data was used to increase coverage whereas spike-in normalised UMI counts were used for burst size inference as it requires accurate expression estimates. Earth’s Mover Distance (EMD) was calculated for 2d kernel density estimates (MASS::kde2d) using the emdist package. To compare dosage compensation between autosomes and the X-chromosome, expression ratios were calculated relative to median of euploid autosomes per cell, keeping only expressed genes (TPM>1) and excluding genes known to escape X-inactivation (list of genes from previous compilation^15^). Next, weighted medians were calculated for chrX, autosomes and individual aneuploid chromosomes/segments where the weights were the number of genes per chromosome for each group.

### Protemic analyses

#### Sample collection

Cell pellets of approximately 1 million cells were collected at 300g for 5 min and washed with ice-cold PBS 5 times to eliminate serum-containing media. Cell pellets were solubilised in 20 μl of 8M urea in 50 mM Tris-HCl, pH 8.5, sonicated in water bath for 5 min before 10 μl of 1% ProteaseMAX surfactant (Promega) in 10% acetonitrile (ACN) and Tris-HCl as well as 1 μl of 100x protease inhibitor cocktail (Roche) was added. The samples were then sonicated using VibraCell probe (Sonics & Materials, Inc.) for 40 sec with pulse 2-2 s (on/off) at 20% amplitude. Protein concentration was determined by BCA assay (Pierce) and a volume corresponding to 25 µg of protein of each sample was taken and supplemented with Tris-HCl buffer up to 90 μl. Proteins were reduced with 3.5 μl of 250 mM dithiothreitol in Tris-HCl buffer, incubated at 37°C for 45 min and then alkylated with 5 μl of 500 mM iodoroacetamide at room temperature (RT) in dark for 30 min. Then 0.5 µg of sequencing grade modified trypsin (Promega) was added to the samples and incubated for 16 h at 37°C. The digestion was stopped with 5 μl cc. formic acid (FA), incubating the solutions at RT for 5 min. The sample was cleaned on a C18 Hypersep plate with 40 μl bed volume (Thermo Fisher Scientific), dried using a vacuum concentrator (Eppendorf). Peptides, equivalent of 25 µg protein, were dissolved in 70 μl of 50 mM triethylammonium bicarbonate (TEAB), pH 7.1 and labelled with TMTpro mass tag reagent kit (Thermo Fisher Scientific) adding 100 µg reagent in 30 μl anhydrous ACN in a scrambled order and incubated at RT for 2 h. The reaction was stopped by addition of hydroxylamine to a concentration of 0.5% and incubation at RT for 15 min before samples were combined and cleaned on a C-18 HyperSep plate with 40 μl bed volume. The combined TMT-labelled biological replicates were fractionated by high-pH reversed-phase after dissolving in 50 μl of 20 mM ammonium hydroxide and were loaded onto an Acquity bridged ethyl hybrid C18 UPLC column (2.1 mm inner diameter × 150 mm, 1.7 μm particle size, Waters), and profiled with a linear gradient of 5–60% 20 mM ammonium hydroxide in ACN (pH 9.0) over 48 min, at a flow rate of 200 μL/min. The chromatographic performance was monitored with a UV detector (Ultimate 3000 UPLC, Thermo Scientific) at 214 nm. Fractions were collected at 30 sec intervals into a 96-well plate and combined in 12 samples concatenating 8-8 fractions representing peak peptide elution.

#### Liquid Chromatography-Tandem Mass Spectrometry Data Acquisition

The peptide fractions in solvent A (0.1% FA in 2% ACN) were separated on a 50 cm long EASY-Spray C18 column (Thermo Fisher Scientific) connected to an Ultimate 3000 nano-HPLC (ThermoFisher Scientific) using a gradient from 2-26% of solvent B (98% AcN, 0.1% FA) in 90 min and up to 95% of solvent B in 5 min at a flow rate of 300 nL/min. Mass spectra were acquired on a Orbitrap Fusion Lumos tribrid mass spectrometer (Thermo Fisher Scientific) ranging from m/z 375 to 1500 at a resolution of R=120,000 (at m/z 200) targeting 4×10^5^ ions for maximum injection time of 50 ms, followed by data-dependent higher-energy collisional dissociation (HCD) fragmentations of precursor ions with a charge state 2+ to 6+, using 45 sec dynamic exclusion. The tandem mass spectra of the top precursor ions were acquired in 3 sec cycle time with a resolution of R=50,000, targeting 1×10^5^ ions for maximum injection time of 150 ms, setting quadrupole isolation width to 0.7 Th and normalised collision energy to 35%.

#### Data preprocessing

Acquired raw data files were analysed using Proteome Discoverer v3.0 (Thermo Fisher Scientific) with MS Amanda v2.0 search engine against *Mus musculus* protein database (UniProt). A maximum of two missed cleavage sites were allowed for full tryptic digestion, while setting the precursor and the fragment ion mass tolerance to 10 ppm and 0.02 Da, respectively. Carbamidomethylation of cysteine was specified as a fixed modification. Oxidation on methionine, deamidation of asparagine and glutamine, as well as acetylation of N-termini and TMTpro were set as dynamic modifications. Initial search results were filtered with 1% FDR using the Percolator node in Proteome Discoverer. Quantification was based on the reporter ion intensities.

#### Data Analysis

Proteomics data from clonal cell populations were provided as normalised protein abundance values per gene per clone. To investigate the relationship between transcript and protein levels, correlation analyses were performed using Xpress-seq spike-in normalised UMI counts. Genes with low RNA expression (normalised UMIs ≤ 1) and proteins with abundance ≤ 1 were excluded. For each clone, a linear regression model was fitted between transcript and protein abundances on a log– log scale and Spearman and correlation coefficients and associated p-values were calculated. Protein abundance ratios were then integrated with gene copy number information to explore the relationship between gene dosage and protein expression; proteins lacking CNV information were excluded from downstream analyses. To compare RNA-level compensation, normalised UMIs were summarised for all genomic segments sharing DNA copy number per cell using 20% trimmed means. Next, summarised values were normalised relative to the median of euploid chromosomes per clone and median per DNA copy and ploidy was calculated and compared relative to protein medians. To calculate observed/expected protein abundance, observed ratios were divided by the expected ratio based on DNA information (DNA_copies/ploidy). Linear models were fitted for log2 observed/expected protein ratios and log2 relative DNA copies and values and 95% confidence intervals up to 2-fold increases (log2 = 1) were predicted from the model.

### Reanalysis of publicly available data

#### Allele-resolved scRNA-seq data

Allelic expression for Smart-seq2 data was obtained^35^ and inference of kinetics was performed as described for Xpress-seq data but using only TPM values.

#### Multi-modal DNTR-seq data

Gene expression and binned DNA count data was obtained^33^ and filtering and ERCC spike-in normalisation was performed using scater/scran as described for Xpress-seq. DNA-level data was GC-corrected (but not mappability-corrected due to already mappability-corrected variable genomic bins) as described in HMMcopy and then segmented as described for DNA-seq. Median segment copies were used for hierarchical agglomerative clustering (using complete agglomeration method on euclidean distances) and trees were cut at a height of 60. Genes in segments containing less than 10 or 50 genes were excluded from cluster-level or global analyses, respectively.

#### TCGA data

Quantified gene expression and gene-level and masked genomic segmented copy number data was obtained for primary tumours and normal solid tissue samples from the TCGA-COAD project id. Genes with gene-level copy numbers between 1-4 and an average TPM across samples >1 were kept for analysis.

### Statistics and data visualisation

All statistical tests were performed in R (v.4.4.2; R Core Team 2021) as two-tailed unless otherwise stated. Data visualisation was generated using the R/ggplot2 package.

## Acknowledgements

This study was supported by grants from the Knut & Allice Wallenberg Foundation (2021.0142 and 2022.0146), the Swedish Research Council (2022-01620), the Swedish Society for Medical Research (CG-22-0260) to BR, and the Swedish Society for Medical Research (PD20-0217) and KAW & SciLifeLab (KAW 2024.0159) to A.L. We thank Vincent Pasque useful discussions in which both parties shared ideas and experimental results before release of our respective studies. Protein identification and quantification were carried out by the Proteomics Biomedicum core facility, Karolinska Institutet (https://ki.se/en/research/proteomics-biomedicum-core-facility). The results shown here are in part based upon data generated by the TCGA Research Network: https://www.cancer.gov/tcga.

## Author contributions

N.P performed all wet lab experiments, analysed data, and wrote the manuscript. S.W. performed DNA-seq and proteomics analyses, curated data, and edited the manuscript. G.M. analysed proteomics data. A.L performed analyses of minibulk SS2, DNA-seq, Xpress-seq and published data, conceived and supervised the study, and wrote the manuscript. B.R. conceived and supervised the study, provided funding, and wrote the manuscript.

## Code availability

Code to reproduce this work is available at Github (https://github.com/reiniuslab/Autosomal_Dosage_Compensation).

## Data availability

Raw and processed sequencing data will be made available upon peer review of this study.

---

**Supplementary Figure 1.**
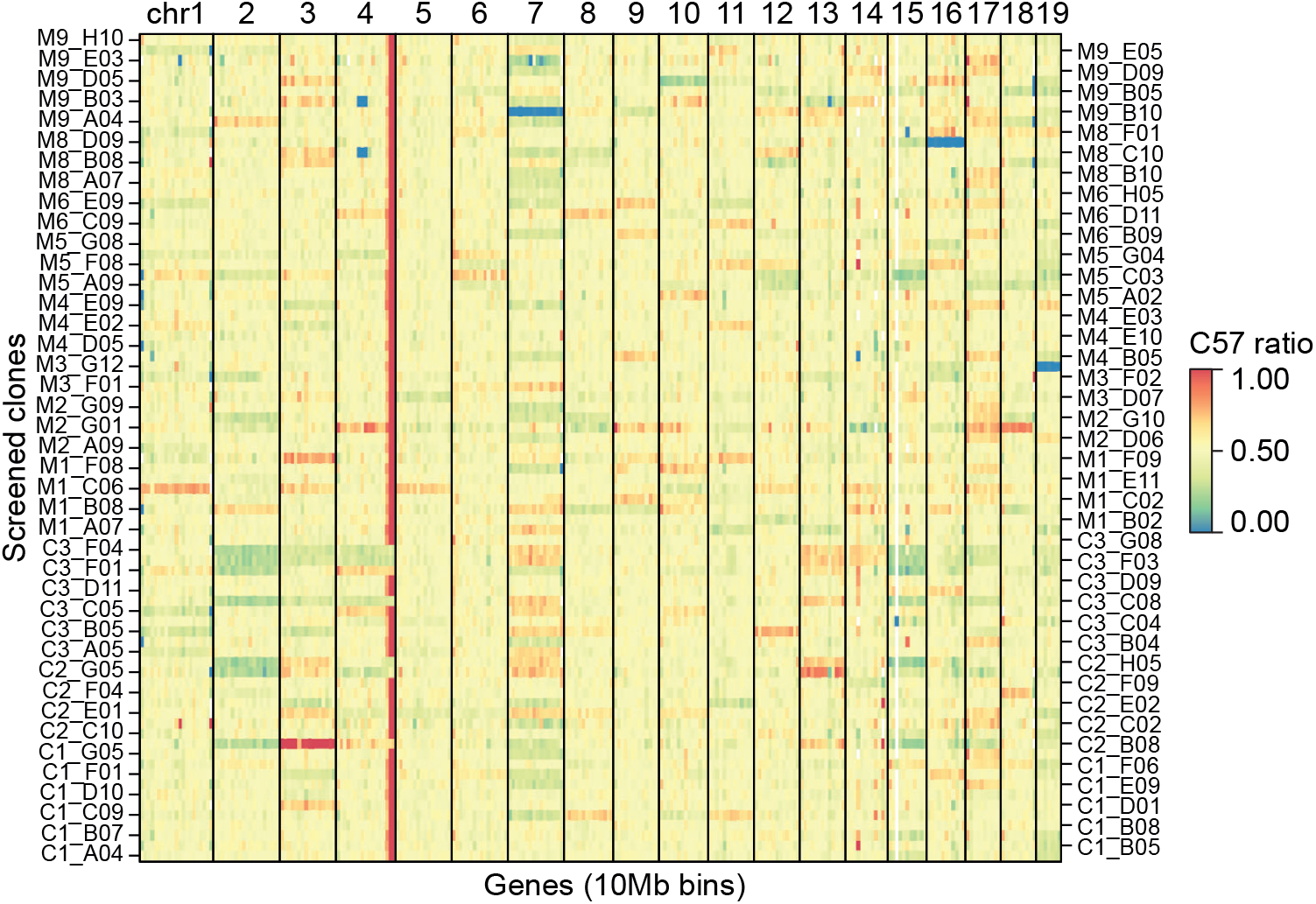
Clonally expanded fibroblasts display varying degrees of aneuploidy. Heatmap of allelic gene expression ratios based on minibulk Smart-seq2 data shown for screened clonal F1 fibroblast lines per autosomal chromosome, divided into 10Mb exression bins. Untreated clones are labelled “C”, and Mitomycin-C treated clones are labelled “M”.

**Supplementary Figure 2.**
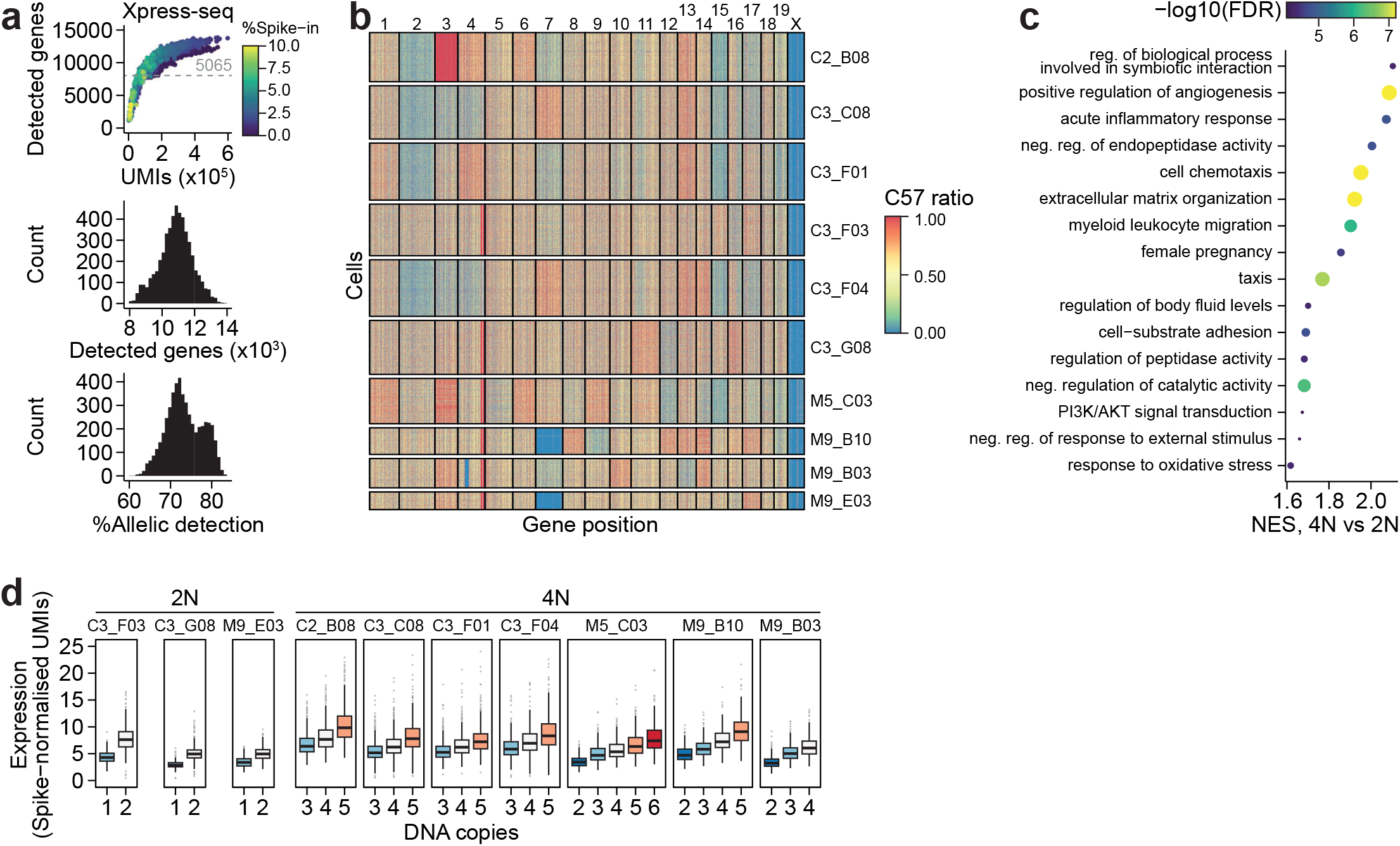
Single-cell RNA-seq on aneuploid clonal lineages. **a**, Quality control of UMI spike-in scRNA-seq (Xpress-seq) aneuploid clonal libraries. Top: Number of genes detected versus number of sequencing reads per cell (n=5,065 cells). Cells with low gene detection (gray dashed line) were excluded from downstream analyses. Middle: Histogram of the distribution of counts on y-axis versus number of genes detected (in thousands) on the x-axis based on Xpress-seq scRNA-seq data. Bottom: Histogram of the distribution of counts on the y-axis versus percentage of allelic detection on the x-axis based on Xpress-seq scRNA-seq data. **b**, Heatmap of allelic gene expression ratios for selected clonal lines used for scRNA-seq experiments, with every row corresponding to an individual cell and every column corresponding to an individual gene, organised by gene position on the chromosome. **c**, Top Gene Ontology terms ranked by enrichment score (–log10 FDR adjusted p-value) for differentially expressed genes between diploid (2N) and tetraploid (4N) clones. d, Boxplots of spike-in normalized UMI gene expression based on Xpress-seq scRNA-seq for selected F1 clonal lines shown in relation to the number of DNA copies in each clone. Number of genes used for each boxplot in this plot. Data shown as median, first and third quartiles, and 1.5x interquartile range.

**Supplementary Figure 3.**
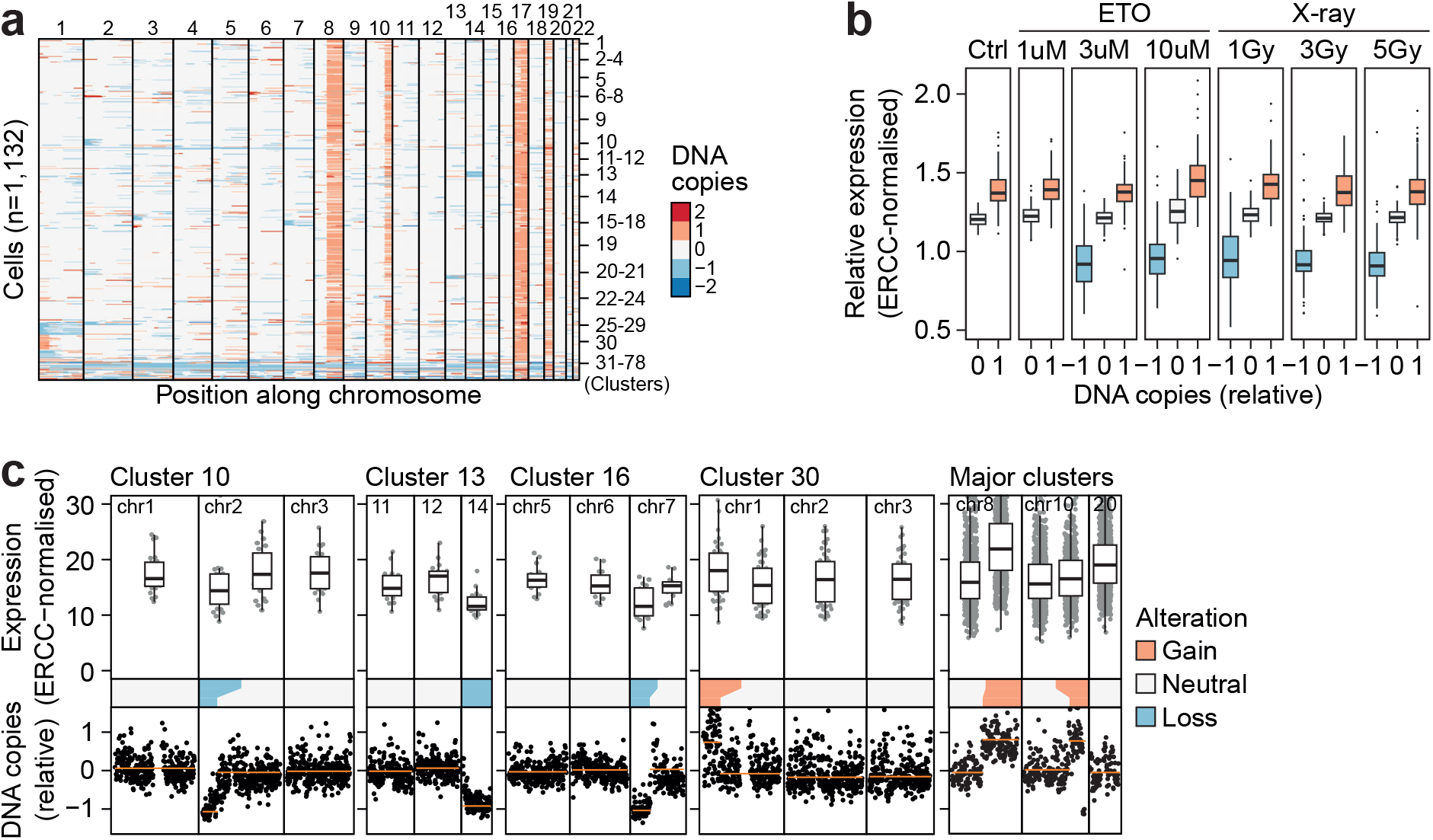
Human adenocarcinoma cell lines (HCT116) display autosomal dosage compensation. **a**, Heatmap of DNA copies in aneuploid human HCT116 cells treated with etoposide or X-rays based on DNTR-seq data with each row corresponding to a cell and each column corresponding to a genomic position, divided into individual chromosomes. Cells were clustered based on shared genetic lesions. The cluster each cell corresponds to is shown on the right. **b**, Boxplots of ERCC-normalised gene expression levels relative to diploid cells shown for each treatment and number of DNA copies. Data shown as median, first and third quartiles, and 1.5x interquartile range. **c**, Top: Boxplots of ERCC-normalised gene expression levels for selected euploid and aneuploid chromosomes for different human adenocarcinoma HCT116 clonal cell clusters. Data shown as median, first and third quartiles, and 1.5x interquartile range. Bottom: Scatterplots of DNA copies corresponding to each of the selected chromosomes and clonal clusters, based on DNTR-seq data.

**Supplementary Figure 4.**
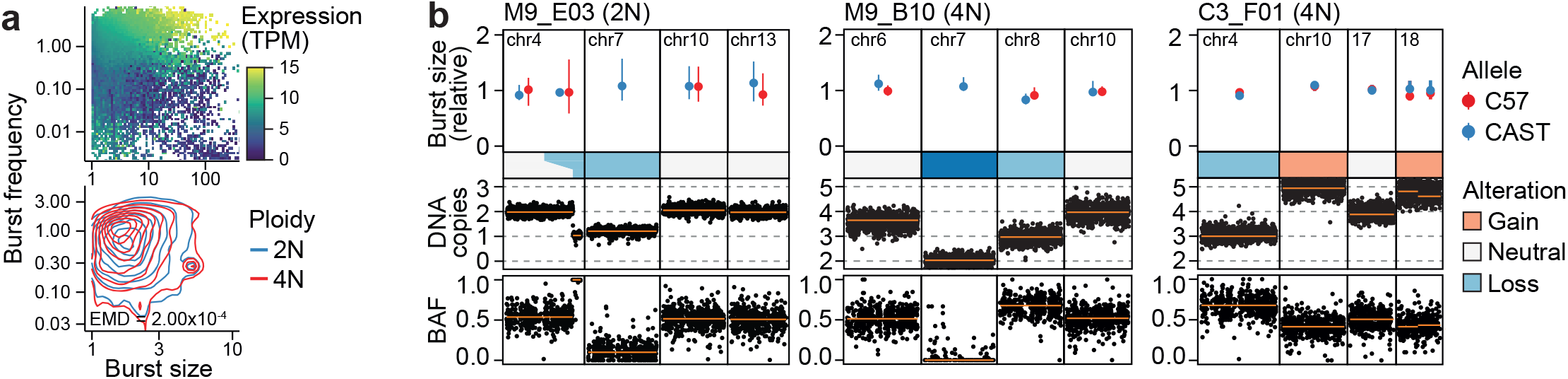
Transcription burst kinetics of aneuploid autosomal chromosomes. **a**, Top: Scatterplot of transcription burst frequency on the y-axis versus transcription burst size on the x-axis based on Xpress-seq data from selected F1 clonal lines. Coloured by gradient of expression level (in TPM). Bottom: 2D kernel density plot of transcription bursting parameters, with burst size represented on the x-axis and burst frequency on the y-axis for diploid (2N) in blue and tetraploid (4N) in red aneuploid clones based on Xpress-seq data. Distance metric is Earth’s mover distance (EMD). Bb, Top: Median transcription burst size, relative to transcription burst size of non-aneuploid chromosomes, shown per allele (C57BL6/J in red and CAST/Eij in blue) for selected diploid (2N) and tetraploid (4N) clones and for specific chromosomes per clone, shown as median and 95% confidence intervals. Middle: Scatterplot of DNA copy numbers per chromosome and clone based on DNA-sequencing data. Bottom: Allele frequency per chromosome and clone based on DNA-seq data.

**Supplementary Figure 5.**
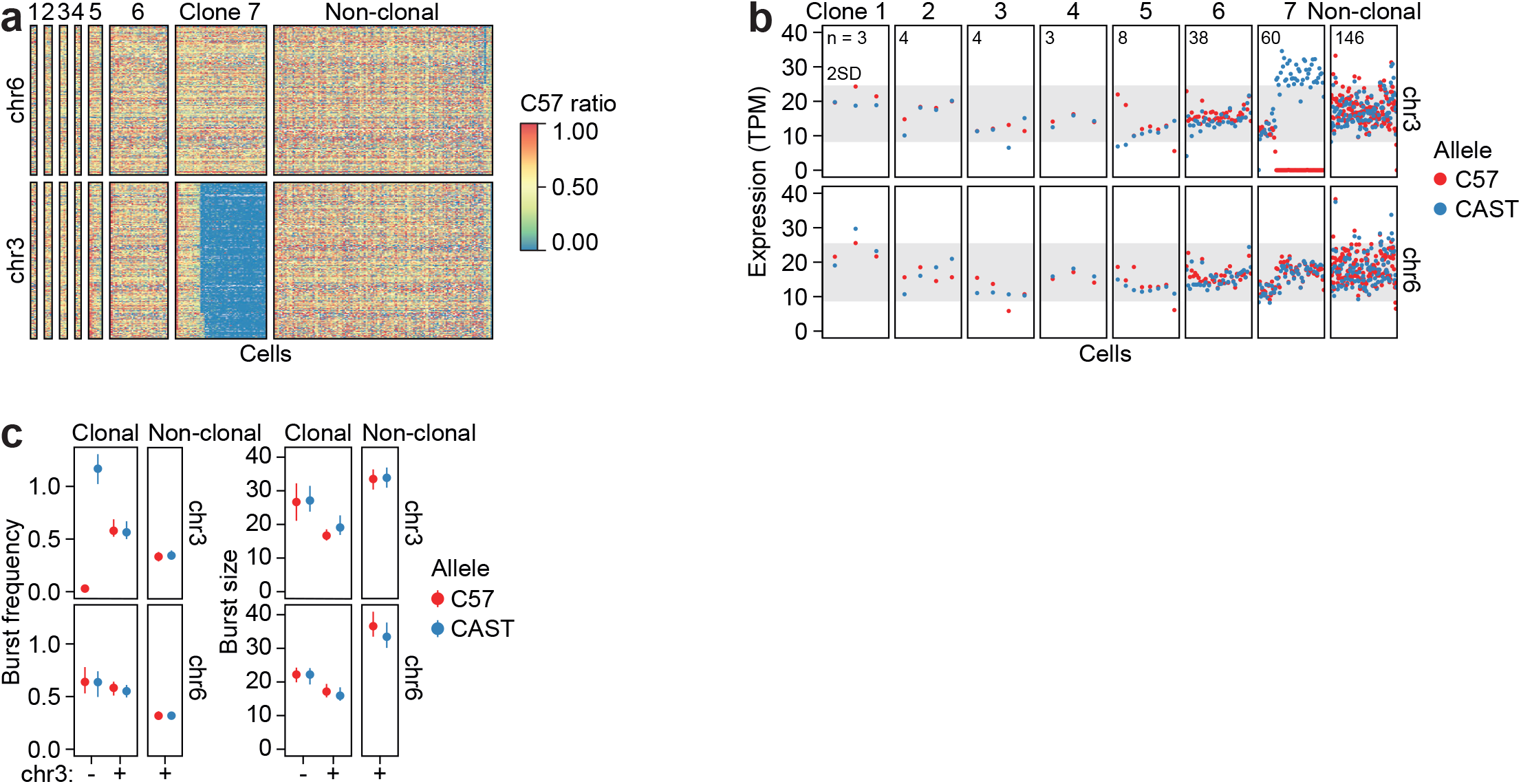
Confirmation of increased burst frequency in an independent aneuploid clonal fibroblast line. **a**, Heatmap of allelic ratios of gene expression based on Smart-seq2 data from an independent dataset, shown for clones 1-7 and a non-clonal fibroblast population. Each row corresponds to a gene located either on chromosome 6 or chromosome 3 and each column corresponds to a cell. **b**, Allele-resolved chromosome expression levels (in TPM) shown per cell for clonal fibroblast lines 1-7 as well as a non-clonal fibroblast population, for chromosomes 3 and 6. C57BL6/J alleles shown in red and CAST/Eij alleles shown in blue. Number of cells per clonal line shown at the top left corner of each individual clonal panel. **c**, Median burst frequency (left panel) and burst size (right size) shown for clonal and non-clonal cells for chromosomes 3 and 6, coloured by allele. C57BL6/J allele shown in red and CAST/Eij allele shown in blue. The x-axis denotes the number of copies for chromosome 3 with ‘-’ corresponding to chr3 monosomy and ‘+’ to disomy (euploidy). Data shown as median and 95% confidence intervals.

**Supplementary Figure 6.**
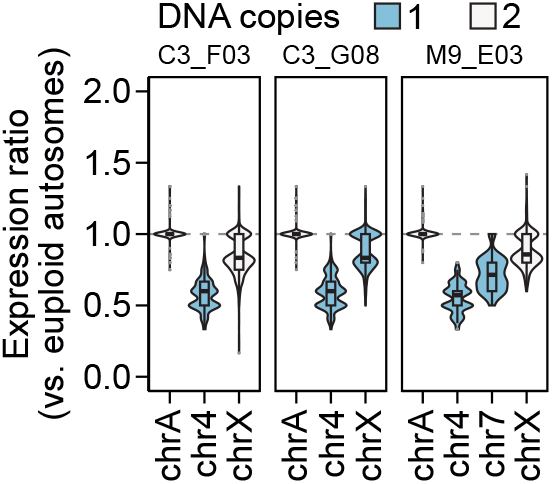
Comparing the magnitude of autosomal upregulation with X-chromosome upregulation. Violin and boxplots of gene expression ratios for euploid and aneuploid autosomes and the X chromosome relative to euploid chromosomes for selected clones based on Xpress-seq data, coloured by number of DNA copies of respective chromosomes. Chromosomal loss resulting in presence of a single copy of a specific chromosome shown in blue, and euploid chromosomes shown in white. Data shown as median, first and third quartiles, and 1.5x interquartile range.

**Supplementary Figure 7.**
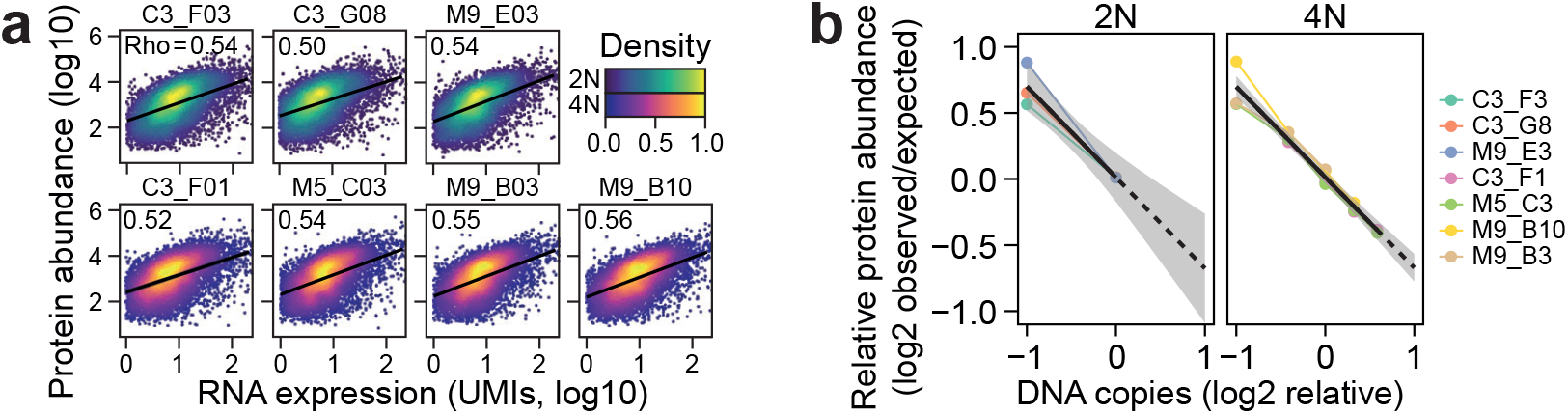
Chromosomal gains and losses are linearly compensated. **a**, Correlation scatterplots of gene expression based on Xpress-seq pseudobulk RNA-seq data (x-axis) and mass spectrometry protein abundance measurements (y-axis) of selected aneuploid clones, coloured by density for diploid (2N) and tetraploid (4N) aneuploid clones. Spearman’s Rho shown at the topleft corner of each individual panel. **b**, Correlation of relative protein abundance (in log2 observed/expected ratio) on the y-axis based on mass spectrometry protein abundance measurements and number of DNA copies (in log2) based on DNA-seq on the x-axis for diploid (2N) and tetraploid (4N) clones, coloured by clone.

